# AtPIG-S, a predicted Glycosylphosphatidylinositol Transamidase Subunit, is critical for pollen tube growth in Arabidopsis

**DOI:** 10.1101/2020.05.06.066910

**Authors:** Nick Desnoyer, Greg Howard, Emma Jong, Ravishankar Palanivelu

**Author notes:** Department of Plant and Microbial Biology, University of Zurich, Zollikerstrasse 107, CH-8008 Zurich Switzerland.

## Abstract

**Background:** Glycosylphosphatidylinositol (GPI) addition is one of the several post-translational modifications to proteins that increase their affinity for membranes. In eukaryotes, the GPI transamidase complex (GPI-T) catalyzes the attachment of pre-assembled GPI anchors to GPI-anchored proteins (GAPs) through a transamidation reaction. A mutation in *AtGPI8* (*gpi8-2*), the putative catalytic subunit of GPI-T in Arabidopsis, is transmitted normally through the female gametophyte (FG), indicating the FG tolerates loss of GPI transamidation. In contrast, *gpi8-2* almost completely abolishes male gametophyte (MG) function. Still, the unexpected finding that *gpi8-2* FGs function normally requires further investigation. Additionally, specific developmental defects in the MG caused by loss of GPI transamidation remain poorly characterized.

**Results:** Here we investigated the effect of loss of *AtPIG-S,* another GPI-T subunit, in both gametophytes. Like *gpi8-*2, we showed that a mutation in *AtPIG-S* (*pigs-1*) disrupted synergid localization of LORELEI (LRE), a putative GAP critical for pollen tube reception by the FG, yet is transmitted normally through the FG. Conversely, *pigs-1* severely impaired male gametophyte (MG) function during pollen tube emergence and growth in the pistil. A *pPIGS:PIGS-GFP* transgene complemented these MG defects and enabled generation of *pigs-1/pigs-1* seedlings, but seemingly failed to rescue the function of AtPIG-S in the sporophyte, as *pigs-1/pigs-1, pPIGS:PIGS-GFP* seedlings died soon after germination.

**Conclusions:** Characterization of *pigs-1* provided further evidence that the FG tolerates loss of GPI transamidation more than the MG and that the MG compared to the FG may be a better haploid system to study the role of GPI-anchoring. *pigs-1* pollen develops normally and thus represent a tool in which GPI anchor biosynthesis and transamidation of GAPs have been uncoupled, offering a potential way to study free GPI in plant development. While previously reported male fertility defects of GPI biosynthesis mutants could have been due either to loss of GPI or GAPs lacking the GPI anchor, our results clarified that the loss of mature GAPs underlie male fertility defects of GPI-deficient pollen grains, as *pigs-1* is defective only in the downstream transamidation step. Our study also provided further evidence that GPI transamidation is essential in seedling development.

## Background

In *Arabidopsis thaliana* (Arabidopsis) plant reproduction, pollen grains land on the stigma of a flower, adhere to the papillae of the stigma, and signal them to initiate hydration and germination of the compatible pollen grain (Mayfield and Preuss, 2000, Wang et al., 2017). Upon germination, a pollen tube will extend through the style and transmitting tract of the pistil and emerge into an ovule chamber, where it targets a female gametophyte (FG) and completes double fertilization. The two synergid cells (synergids) found at the micropylar end of the FG are transfer cells, which are specialized in secretion of factors for proper communication with and reception of the pollen tube (Huang and Russell, 1992).

Synergids develop a region of highly dense cell wall and plasma membrane invaginations at their micropylar end called the filiform apparatus (FA) that is shared by both synergids. Extensive plasma membrane invaginations in the FA increases the total surface area of the synergids, promoting diffusion of proteins and molecules out of the cell and generating an extracellular microenvironment conducive to attraction (Higashiyama and Takeuchi, 2015) and reception (Qu et al., 2015) of pollen tubes. After entering the ovary chamber, pollen tubes respond to the chemoattractants from the synergid cells, migrate steadfastly to an ovule, enter the micropylar opening in the ovule, and reach the FA of the synergid cells, the first physical point of contact between the pollen tube and the synergid cells (Johnson et al., 2019). The cessation of pollen tube growth in one of the synergids (receptive synergid cell) and subsequent release of sperm cells to induce double fertilization is collectively called pollen tube reception, a process that is regulated by genes expressed in both gametophytes (Johnson et al., 2019).

In the FG, LORELEI (LRE), a predicted glycosylphosphatidylinositol (GPI)-anchored protein (GAP), binds the receptor-like kinase FERONIA (FER) to form a co-receptor complex critical for the perception of an unknown ligand to transduce downstream signaling necessary for pollen tube reception (Liu et al., 2016, Li et al., 2015). Homozygous *lre-7/lre-7* (Tsukamoto et al., 2010), *fer-4* mutant (Duan et al., 2014), and *lre-7, fer-4* double mutant pistils (Liu et al., 2016) all show similar levels (~80%) of reduction in seed set, indicating both proteins function together in the same pathway. Using a *pLRE:LRE-cYFP* translational fusion construct, we previously showed that the seed set defect in *lre* mutants can be fully rescued and that LRE-cYFP localizes in the FA of synergid cells (Liu et al., 2016). Consistent with apical sorting expected of a plant GPI-anchored protein (GAP), roughly 68% of all cYFP signal in the synergids expressing LRE-cYFP accumulate in the FA. Deletion of both predicted omega sites in LRE, residues to which GPI is expected to be post-translationally attached, or complete ablation of the GAS region in the C-terminus of LRE protein, dramatically disrupts the localization of LRE-cYFP in the FA. While this disruption in localization of these mutant LRE-cYFP is suggestive of their failure to receive a GPI anchor, both mutants fully complement pollen tube reception and seed set defects in *lre-7/lre-7* (Liu et al., 2016). Even replacing the GAS region in LRE, with the transmembrane domain of FER, fully complements *lre-7* female fertility defects, showing that LRE can embed in the plasma membrane in a completely different form and still function in pollen tube reception to near normal levels (Liu et al., 2016).

To address whether the loss of GPI anchor addition to other FG-expressed GAPs is tolerated, in addition to the *cis*-mutations in LRE, we previously also analyzed a mutation in a subunit of the GPI transamidase (GPI-T), the transacting complex responsible for GPI anchor addition to GAPs by transamidation (Liu et al., 2016). A T-DNA insertion in *GPI8* (*gpi8-2*), the putative catalytic subunit of GPI-T, affected the polar localization of LRE-cYFP in the FA of synergids at a similar level to the *cis*-mutations in GAS region of LRE. Still, pollen tube reception and seed set in *gpi8-2/*+ pistils was indistinguishable from wild-type pistils, and near normal *gpi8-2* transmission through the FG suggested that loss of GPI anchor addition to GAPs in the FG does not disrupt its function. In contrast, transmission of *gpi8-2* through the MG was abolished and *gpi8-2* homozygotes could not be established (Liu et al., 2016). The precise MG function that is affected in *gpi8-2* and confirmation of *AtGPI8* function in MG by complementation assays have not been reported; consequently, which aspect of pollen function in fertilization (pollen development, pollen tube emergence, pollen tube growth, and pollen tube reception) is disrupted from loss of GPI anchor addition to GAPs expressed in the MG remains unknown.

The normal function of *gpi8-2* FGs was rather unexpected, as GPI anchor addition to GAPs is the most complex and metabolically costly lipid post-translational modification in eukaryotes (Orlean and Menon, 2007) and is most likely used as a mechanism for the membrane attachment of many other GAPs in FG function. To obtain additional evidence in support of the finding that loss of GPI anchor addition to GAPs disrupts MG functions more than the FG (Liu et al., 2016), in this study, we investigated the defects in Arabidopsis FG and MG caused by a mutation in *AtPIG-S* (*At3g07180*), a homolog of yeast and human PIG-S and a putative subunit of GPI-T. In addition, we utilized the reporter gene in the T-DNA that disrupted *AtPIG-S* to pinpoint specific MG phenotypes caused by the loss of AtPIG-S in GPI-T.

The GPI-T complex consists of 5 subunits – PIG-K/GPI8, GPAA1/GAA1, PIG-S/GPI17, PIG-T/GPI16, and PIG-U/GAB1 – in mammals and yeast, respectively (Gamage and Hendrickson, 2013). Arabidopsis homologs of each of these GPI-T subunits have been identified (Ellis et al., 2010) and is comprised of 5 subunits - PIG-K/GPI8 (*At1g08750*), GAA1 (*At5g19130*), PIG-S (*At3g07180*), PIG-T (*At3g07140*), and PIG-U (*At1g63110*) (Luschnig & Seifert 2011). Of these, only the catalytic GPI8 subunit had been characterized in greater detail in Arabidopsis (Bundy et al., 2016, Liu et al., 2016). In this study, we showed that a mutation in *AtPIG-S*, a putative GPI-T subunit in Arabidopsis, does not affect the function of the FG even though it disrupts localization of LRE-cYFP. Furthermore, our results demonstrated that *pigs-1* transmission through the MG is almost completely abolished due to defects at the stages of pollen tube emergence and tube growth, revealing the importance of GPI anchor addition to the MG-expressed GAPs during pollen pistil-interactions. Our results reported in this study using *pigs-1* are consistent with previous results obtained using *gpi8-2* and showed that the FG is far more tolerant than the MG to the loss of GPI-T function and GPI anchor addition to GAPs.

Characterization of the *pigs-1* mutation also showed that GPI transamidation-deficient pollen grains develop normally and thus represent a tool in which GPI anchor biosynthesis and GPI anchor addition to GAPs have been uncoupled, offering a potential way to study of free GPI in plant development. Restoration of normal tube growth in *pigs-1* pollen by supplying a wildtype copy of the *AtPIG-S* fused to *GFP* and expressed from its own promoter (*pPIGS:PIGS-GFP*) allowed the generation of *pigs-1/pigs-1* seeds. However, for reasons that are not well understood, the transgene did not complement AtPIG-S function in the sporophyte, resulting in lethality of *pigs-1/pigs-1, pPIGS:PIGS-GFP* seedlings directly following emergence from the seed coat. Still, lethality of *pigs-1/pigs-1* seedlings demonstrated the importance of AtPIG-S and GPI anchor addition to GAPs in early seedling development.

## RESULTS

### A mutation in *AtPIG-S* disrupted localization of LORELEI in the filiform apparatus of synergid cells

To examine if loss of GPI-T subunits (and by extension, loss of GPI anchoring) disrupts function of the Arabidopsis FG more than the MG, we obtained a T-DNA insertion in *AtPIG-T* (*pigt-1*). PIG-T is the central component of GPI-T and is critical for the formation of complex by binding the luminal domains of GAA1, the largest subunit of GPI-T that likely presents GPI to the complex reaction center, and PIG-S, a peripheral component with unknown function (Ohishi et al., 2001). As was done with *gpi8-2* (Liu et al., 2016), we used disruption of LRE-cYFP localization in the FA of synergids of the *pigt-1* mutant as a marker to characterize AtPIG-T function in the FG. Surprisingly, all ovules in *pigt-1/+, pLRE:LRE-cYFP/ pLRE:LRE-cYFP* pistils showed a localization pattern that is indistinguishable from LRE-cYFP localization in wild-type background (Additional file 1), despite AtPIG-T being predicted to encode a functionally important subunit of GPI-T (Ohishi et al., 2001). RT-PCR and RT-qPCR experiments using 14-day old homozygous *pigt-1/pigt-1* seedlings showed that full-length *AtPIG-T* transcripts were not detected in *pigt-1* seedlings and that truncated *AtPIG-T* mRNA transcripts, consisting of exons 1-4, accumulated to a level that is only 40% compared to wild type (Additional file 2). Based on these results we concluded that *AtPIG-T* transcripts lacking exon 5, which encodes the cytosolic domain and ER-retrieval motif of PIG-T (Fraering et al., 2001), coupled with the lower levels of these partial transcripts, is still not sufficient to cause a noticeable reduction in LRE-cYFP localization in the FA of *pigt-1* synergids. Consequenlty, we did not use *pigt-1* any further to investigate if disrupting GPI-T function affects Arabidopsis MG more than the FG.

We next obtained a T-DNA mutant line (*pigs-*1) in *AtPIG-S*, another putative subunit of GPI-T. Homozygous *pigs-1/pigs-1* could not be obtained (Table 1), so we evaluated if *pigs-1* disrupts *LRE* localization in the FA by analyzing localization of LRE-cYFP in pistils heterozygous for *pigs-1* and homozygous for *pLRE:LRE-cYFP* (Fig. 1). We detected two patterns of LRE-cYFP localization in *pigs-1/*+ pistils. In one portion of ovules (~60%, n=334; Fig. 1e), we detected expected LRE-cYFP localization in the FA and in puncta in the synergid cytoplasm (Fig. 1f), as reported previously (Liu et al., 2016). In the remaining ovules (~40%, n=334; Fig. 1e), LRE-cYFP localization was aberrantly diffused throughout the synergids (Fig. 1g). The two types of localization pattern observed here were similar to ovules in heterozygous *gpi8-2* pistils that we previously reported (Liu et al., 2016). Under similar assay conditions, almost 100% of ovules in wild-type pistils carrying the *pLRE-cYFP* transgene had the expected polarized LRE-cYFP localization pattern in synergids (n=229; Fig. 1e). When averaging the relative integrated signal density within the FA of ovules with diffused localization in a *pigs-1/*+ pistil, they showed about a 50% reduction in signal in the FA and had substantially more signal dispersed towards the chalazal end of the synergid cell compared to that in sibling ovules with a polarized LRE-cYFP localization in the FA (Figs. 1h and i). These results showed that *AtPIG-S* is important for polarized localization of LRE in the FA, likely by mediating GPI anchor addition to LRE-cYFP.

**Figure 1.**
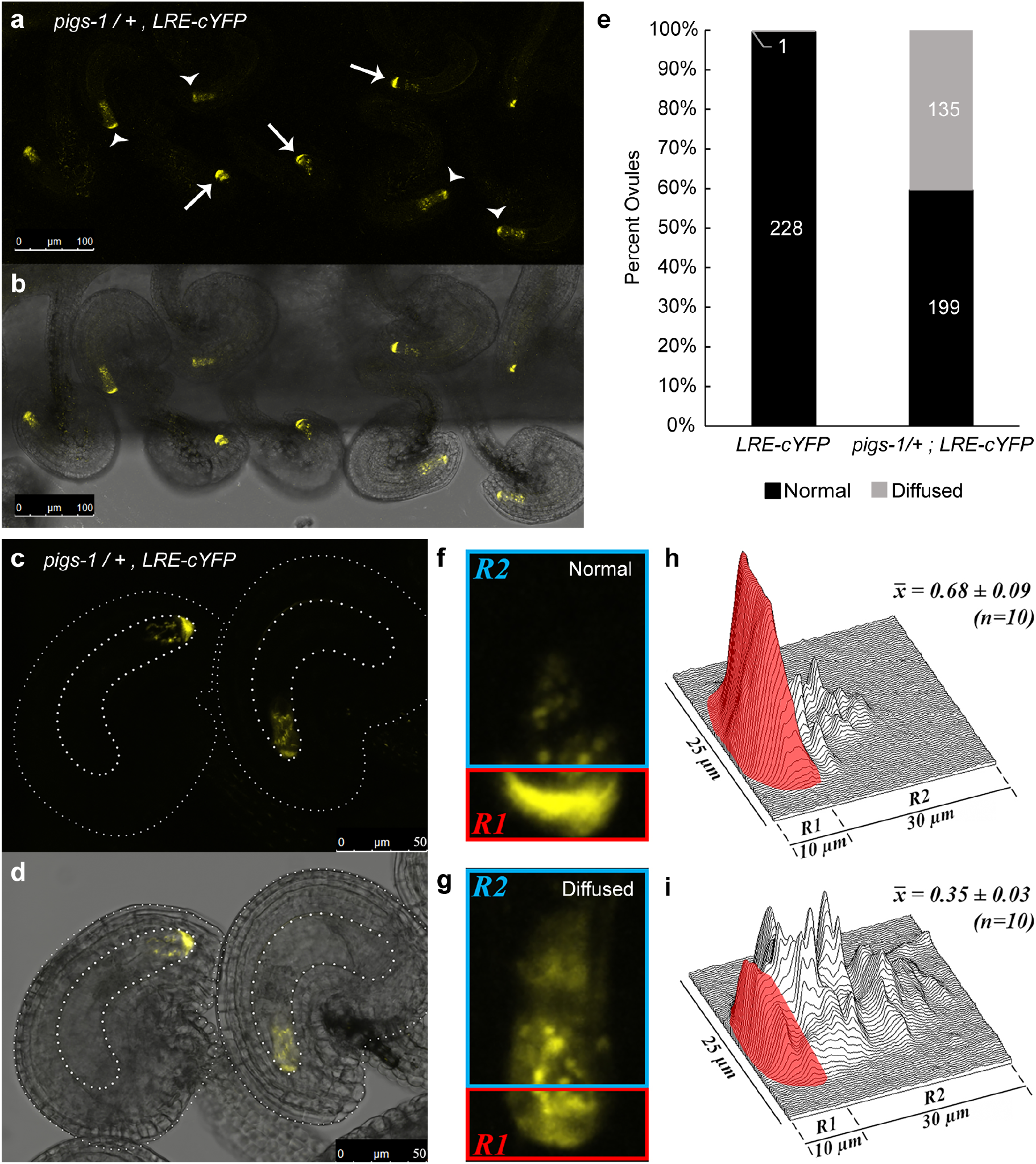
*pigs-1* mutation disrupts polar localization of LRE-cYFP in the filiform apparatus of synergid cells. (**a**-**b**) Localization of the LRE-cYFP fusion protein in a *pigs-1/*+ pistil. (**a**) Fluorescent image of a portion of the pistil captured in the YFP channel of a confocal microscope and (**b**) Merged view of fluorescent image in (**a**) with the bright field image of the same portion of the pistil captured in (**a**). White arrows, FGs with a polarized cYFP localization in the filiform apparatus; white arrowheads, sibling FGs with a diffuse cYFP localization throughout the synergids. Bar = 100 μm. (**c**-**d**) Enlarged view of two ovules within a *pigs-1*/+ pistil Bar = 50 μm. (**c**) Fluorescent image captured in the YFP channel of a confocal microscope and (**d**) Merged view of fluorescent image in (**c**) with the bright field image of the same ovules captured in (**c**). (**e**) Percent ovules showing normal and diffuse LRE-cYFP localization in synergid cells of wild-type and *pigs-1*/+ pistils. Numbers in the column refer to total number of ovules scored for indicated categories. (**f**-**g**) Representative images of (**f**) normal and (**g**) diffuse LRE-cYFP localization patterns scored within synergid cells of a *pigs-1/*+ pistil. (**h**-**i**) Surface plots of (**h**) normal and (**i**) diffuse LRE-cYFP fluorrescent signal scored within synergid cells of a *pigs-1/*+ pistil. X̅ values are average raw integrated densities of cYFP signal in filiform apparatus portion of the synergid cell boxed in red (R1) divided by total signal in R1+R2, where R2 represents remainder of the synergid cell boxed in blue.

**Table 1.** Reduced transmission of the *pigs-1* and *gpi8-2* mutations through the male gametophyte.

| Female parent | Male parent | Observed No. of progeny | | TE<br>(R/S) | $\chi^2$ | P-value |
| --- | --- | --- | --- | --- | --- | --- |
|  |  | Basta<sup>R</sup>* | Basta<sup>S</sup>* |  |  |  |
| *BastaR/+* | Wild type | 120 | 131 | 0.92 | 0.48 | 0.49 |
| *lre-7/+* | Wild type | 29 | 181 | 0.16 | 110 | <0.001 |
| *gpi8-2/+* | Wild type | 177 | 169 | 1.05 | 0.19 | 0.67 |
| *pigs-1/+* | Wild type | 255 | 268 | 0.95 | 0.32 | 0.57 |
| Wild type | *BastaR/+* | 189 | 193 | 0.98 | 0.04 | 0.84 |
| Wild type | *gpi8-2/+* | 0 | 607 | 0 | 607 | <0.001 |
| Wild type | *pigs-1/+* | 8 | 483 | 0.02 | 460 | <0.001 |
| Wild type | *pigs-1/+; PIGS-GFP* | 108 | 142 | 0.76 | 4.62 | 0.03 |

Wild type, Columbia-0 ecotype
\*Basta resistant (Basta<sup>R</sup>) and Basta susceptible (Basta<sup>S</sup>) progeny. Basta resistance gene is linked with the T-DNA that is inserted into the *AtGPI8* (*gpi8-2*) (Liu et al., 2016) and *LRE* (*lre-7*), *AtPIGS* (*pigs-1*), and in an unknown location in the genome of the *BastaR/+* line (Johnson et al., 2004).
TE, Transmission efficiency was calculated as the quotient of the number of basta resistant (R) divided by basta susceptible (S) progeny of the indicated cross.
$\chi^2$ was calculated based on the expectation of a 1:1 segregation of basta resistance to susceptibility in the progeny. *PIGS-GFP* refers to the T2 line of *pPIGS:PIGS-GFP* #15-3.

### Transmission of *pigs-1* is primarily affected in the male gametophyte owing to defects in pollen tube emergence and growth

Disruption of LRE-cYFP localization in the FA of ~40% of ovules in a *pigs-1/*+ pistil coupled with our inability to identify a *pigs-1* homozygote suggested that AtPIG-S may play an important role in the FG, such as affecting GPI anchor addition to LRE and other synergid-expressed GAPs. Additionally, it could also play an important role in the MG. We therefore investigated the role of *AtPIG-S* in either of the two gametophytes by performing reciprocal crosses to analyze the transmission efficiency (TE) of the *pigs-1* mutation through the gamteophytes using the basta resistance marker linked to the *pigs-1* T-DNA (Table 1 and Fig. 2a).

**Figure 2.**
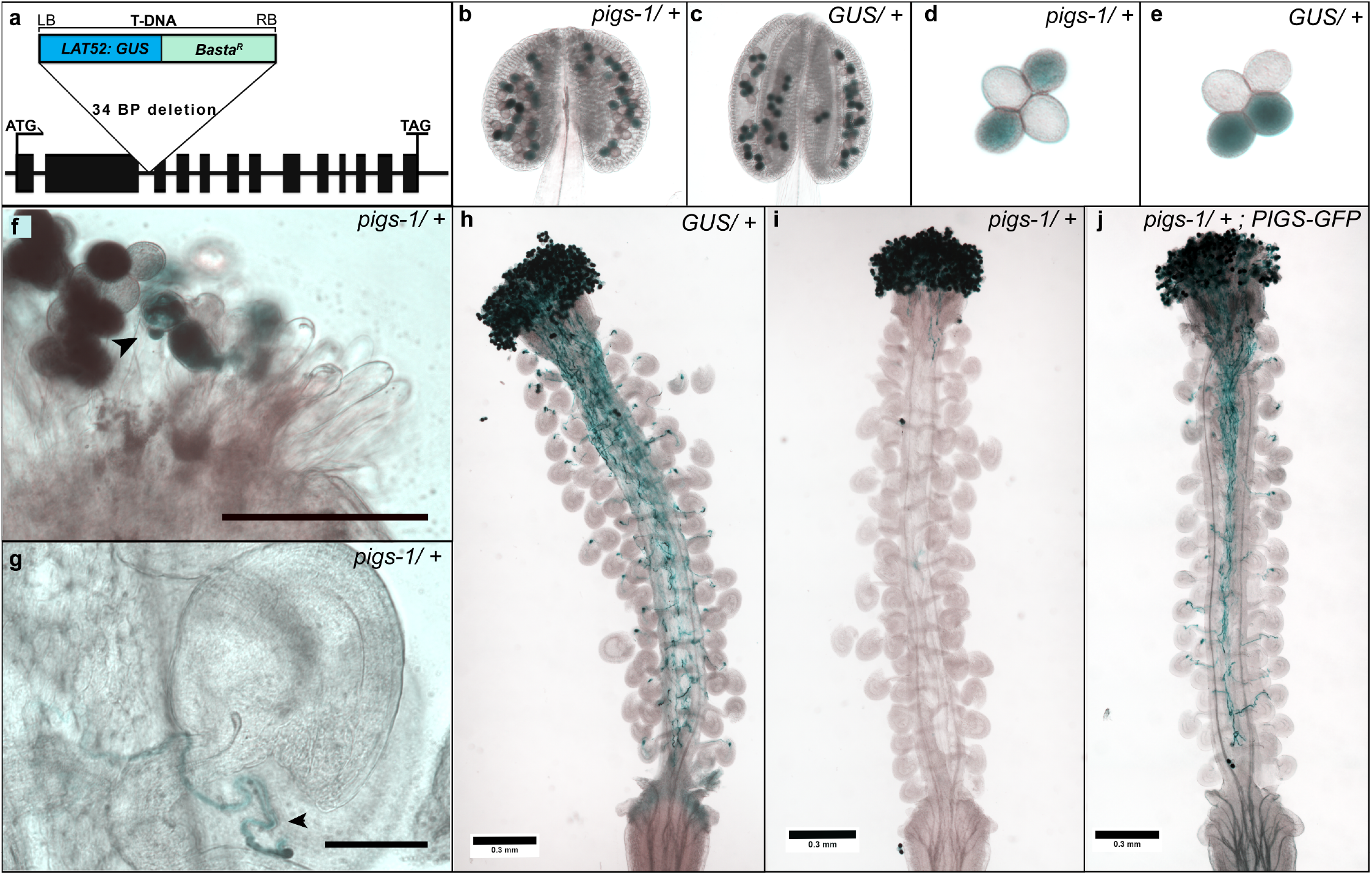
*pigs-1* mutation disrupts pollen tube emergence and growth *in vivo*. (**a**) Gene Model of *AtPIG-S* (*AT3G07180*) with a T-DNA insertion (*SAIL_162_D06*) carrying the basta resistance and *pLAT52:GUS* genes. Black rectangles indicate exons and lines indicate introns and UTRs. Basta resistance gene is under the control of the *35S* promoter and a post meiotic, pollen-specific promoter of *LAT52* drives the beta-glucronidase reporter (*GUS*) gene. Both junctions of T-DNA insertion sites in the *AtPIGS* gene were sequenced to identify the insertion point in the second intron, which caused a 34bp deletion. (**b**-**c**) Anther locules containing indehiscent pollen tetrads in *pigs-1/*+ (**b**) or wild-type backgrounds (**c**) were stained with X-Gluc for GUS activity. (**d**-**e**) Released pollen tetrads from (**d**) *pigs*-1*/*+ or (**e**) wild-type anthers show indistinguishable morphology and *GUS* expression (n=100, in each genotype). (**f**) Stigma of wild-type pistil pollinated with *pigs-1/*+ stained for GUS activity 18 hours after pollination (HAP). Black arrowhead, *pigs-1* pollen showing pollen tube emergence defect. Scale bar 100 uM. (**g**) Rare instance of *pigs-1* pollen tube approaching wild-type ovule within a wild-type pistil pollinated with *pigs-1*/+ and stained for GUS activity 18 HAP. *pigs-1* pollen tube grew along the funiculus and failed to target the micropyle. Black arrowhead, *pigs-1* pollen tube growing away from the micropyle. Scale bar 100 uM. (**h**-**j**) Light micrograph of wild-type pistil pollinated with either (**h**) *pLAT52:GUS/*+, (**i**) *pigs-1/*+, or (**j**) *pigs-1/*+ transformed with *pPIGS:PIGS-GFP* line #15-3 pollen and stained for GUS activity 18 HAP. Scale bars 0.3 mm.

Despite the disrupted LRE-cYFP localization in *pigs-1+* pistils, the segregation ratio of basta-resistant seeds collected from *pigs-1/*+ pistils that were pollinated with wild-type pollen indicated that there is normal TE of *pigs-1* through the FG (95%; Table 1). In contrast, only 2% of seeds collected from wild-type pistils pollinated with a *pigs-1/*+ pollen were basta-resistant, revealing an almost complete abolishment of *pigs-1* transmission through the MG (Table 1). Furthermore, the transmission of *pigs-1* through the MG could not be restored by limited pollination of *pigs-1/*+ pollen on wild-type pistils (Table 2), indicating that the decreased transmission of *pigs-1* through the MG is not attributable to mutant pollen being outcompeted by wild-type pollen; instead, these results point to inherent defects in *pigs-1* pollen function.

**Table 2.** Limited pollination does not restore male transmission in *pigs-1* or *gpi8-2*.

| Female parent | Male parent | Observed No. of progeny | | TE<br>(R/S) | $\chi^2$ | P-value |
| --- | --- | --- | --- | --- | --- | --- |
|  |  | Basta<sup>R</sup>* | Basta<sup>S</sup>* |  |  |  |
| Wild type | *pigs-1/+* | 3 | 121 | 0.02 | 112.3 | <0.001 |
| Wild type | *gpi8-2/+* | 1 | 150 | 0.01 | 147.0 | <0.001 |
| *pigs-1/+* | Wild type | 51 | 71 | 0.72 | 3.28 | 0.07 |
| *gpi8-2/+* | Wild type | 47 | 63 | 0.75 | 2.33 | 0.13 |

\*Basta herbicide resistant (Basta<sup>R</sup>) or susceptible (Basta<sup>S</sup>) progeny. Basta resistance gene is linked with the T-DNA that is inserted into the *GPI8* or *PIGS* genes (*pigs-1* and *gpi8-2* mutants, respectively).
TE, Transmission efficiency was calculated as the quotient of basta resistant (R) divided by susceptible (S) progeny of the indicated cross in which only a limited number of pollen grains (<40) were used.
$\chi^2$ was calculated based on the expectation of a 1:1 segregation of basta resistance to susceptibility in the progeny.

To identify the defect underlying the reduced transmission of *pigs-1* through the male gametophyte, we utilized the *pLAT52:GUS* gene in the *pigs-1* T-DNA as a reporter to assess pollen development and function (Fig. 2a). The *pigs-1* mutation is in a *quartet/quartet* background that prevents microspores from separating even after pollen maturation (Preuss et al., 1994), consequently all four products of every meiosis are preserved, allowing direct comparison of the wild-type and mutant pollen. Indehiscent and dehiscent pollen tetrads from *pigs-1*/+ plants were stained for GUS activity (n = 100) and in both instances, *pigs-1* pollen grains (GUS-positive) had a morphology that is indistinguishable from the wild-type pollen (GUS-negative) on the same tetrad and when compared to all pollen grains in the control line (*pLAT52:GUS/*+ tetrads without any *pigs-1* mutation) (Figs. 2b-e), indicating that *pigs-1* pollen grains have undergone normal development. In addition, *pigs-1* pollen grains expressed *GUS* similar to *pLAT52:GUS* pollen tetrads in wild-type background, indicating that mature *pigs-1* pollen grains are metabolically active and viable.

When *pigs-1/*+ pollen grains were crossed to wild-type pistils, the *pigs-1* pollen showed dramatic pollen tube growth defects. Unlike wild-type pistils pollinated with pollen carrying *pLAT52:GUS* in the wild-type background (Fig. 2h), wild-type pistils pollinated with *pigs-1/*+ pollen and stained for GUS activity showed many non-germinated or barely emerged *pigs-1* pollen tubes (Figs. 2f and i). Many *pigs-1* pollen grains that appeared to have germination defects remained on the stigma despite GUS staining procedure and mounting onto slides, suggesting that these pollen grains had likely successfully adhered to the stigma papillae, a critical first step in compatible pollen-pistil interactions (Johnson et al., 2019). Of the pollen grains that germinated pollen tubes, the average longest *pigs-1* pollen tube in these pistils was 0.38mm ± 0.09 (n=5), whereas wild-type *pLAT52:GUS* pollen tubes grew the entire length of pistils with an average longest length of 2.13mm ± 0.15 (n=5). While mutant *pigs-1* pollen defects primarily manifest at the stage of pollen tube emergence and growth, in one rarely observed instance of a *pigs-1* pollen tube growing along the funiculus of an ovule, disrupted micropylar guidance was observed (Fig. 2g). Based on these results, we concluded that *pigs-1* pollen is defective in pollen tube emergence and growth and that these defects underlie the nearly abolished transmission of *pigs-1* mutation through the pollen.

### The *pPIGS:PIGS-GFP* transgene rescued pollen tube emergence and growth defects in *pigs-1* pollen

To confirm that the observed MG defects in *pigs-1* pollen are solely due to disruption of the *AtPIG-S* gene, we generated a *pPIGS:PIGS-GFP* translational fusion construct in which the 1500bp upstream sequence of the *AtPIG-S* transcription start site drove the expression of GFP fused to the N-terminus of the *AtPIG-S* genomic sequence (Additional file 3a). Unfortunately, PIGS-GFP signal was not detectable in seedlings or pollen grains of T1 plants similar to a previously reported GPI-T subunit fusion construct, AtGPI8-EGFP (Bundy et al., 2016). To verify if the *pPIGS:PIGS-GFP* is expressed, we infiltrated *Agrobacterium* carrying the *pPIGS:PIGS-GFP* construct into *N. benthamiana* leaves and checked if the leaves showed transient expression of PIGS-GFP. We detected GFP signal in the *N. benthamiana* leaves that were infiltrated with *pPIGS:PIGS-GFP* (Additional file 3c and e) but not the leaves that were infiltrated with the helper plasmid only (Additional file 3b and d). Interestingly, GFP signal was not detectable 3 days post infiltration, but substantial GFP signal accumulated by 12 days post infiltration suggesting that while the AtPIG-S-GFP fusion protein is stable, the promoter activity of *pPIGS* used in the construct or incorporation rate of the fusion protein into the GPI-T complex is quite low. Additionaly, consistent with the known residency of the GPI-T complex in the ER, the PIGS-GFP signal in the pavement cells showed a localization pattern that is suggestive of proteins accumulating in the ER (Additional file 3c).

**Table 3.** Seedling lethality of *pigs-1/pigs-1*; *pPIGS:PIGS-GFP* on MS plates.

| Selfed parent | Observed No. of progeny |  | % Lethal |
| --- | --- | --- | --- |
|  | Normal | Lethal |  |
| *pigs-1/+* | 346 | 0 | 0 |
| *pigs-1/+* ; *PIGS-GFP* #15-3 | 315 | 78 | 19.85 |
| *pigs-1/+* ; *PIGS-GFP* #7-25 | 140 | 32 | 18.60 |
| *pigs-1/+* ; *PIGS-GFP* #25-6 | 324 | 127 | 28.16 |

Selfed seeds were placed onto MS plates containing no antibiotic or basta.
Seedling phenotypes were scored according to images and descriptions in Fig. 3.
*PIGS-GFP* refers to three independent transformant T2 lines of *pPIGS:PIGS-GFP*

To examine if the *pPIGS:PIGS-GFP* construct could complement pollen tube emergence and growth defects in *pigs-*1, we analyzed three independent T1 transformant lines in a *pigs-1/*+ background (Lines 7, 15, and 25). In each independent transformant line, we identified T2 individuals that were heterozygous for the *pigs-1* mutation (identified by GUS staining of the tetrads) and carried at least one copy of the *pPIGS:PIGS-GFP* transgene (based on hygromycin resistance linked to the transgene) and used them in complementation experiments. Wild-type pistils were pollinated with pollen from these T2 plants and stained for GUS activity 18 hours after pollination. In all three independent transformant lines, *PIGS-GFP* rescued the pollen tube emergence and growth defects in *pigs-1* pollen, as *pigs-1* pollen tubes (GUS-positive) grew the entire length of the pistils (Fig. 2j). Importantly, the longest pollen tubes in wild-type pistils crossed to pollen from three independent transformants was restored to the full length of the pistil and averaged 2.2 ± 0.16mm, 2.2 ± 0.11, and 2.14mm ± 0.14mm in length, respectively. Furthermore, presence of the *pPIGS:PIGS-GFP* transgene rescued the reduced TE of the *pigs-1* mutation through the MG, as the TE of the *pigs-1* mutation significantly increased from 0.02 without the *pPIGS:PIGS-GFP* transgene to a TE of 0.76 with the *pPIGS:PIGS-GFP* transgene (Table 1). Although *pPIGS:PIGS-GFP* did not generate a detectable GFP signal in the pollen grains or pollen tubes of any of the three independent transformant lines, the successful complementation of pollen tube growth defects indicates that the transgene is likely expressed. Together, these results indicated that the pollen tube emergence and growth defects in *pigs-1* pollen carrying the *pPIGS:PIGS-GFP* transgene were restored to normal levels.

### *pigs-1/pigs-1, pPIGS:PIGS-GFP* seedlings died after emerging from the seed coat

Despite this restoration of pollen tube growth, out of 75 T2 plants from the three independent *pPIGS:PIGS-GFP* transformant lines, we never detected a T2 plant that produced pollen tetrads homozygous for GUS (4:0 GUS+: GUS-tetrad), the marker linked with the *pigs-1* mutation. These results suggested that homozygous *pigs-1* plants are not present or occur at a very low frequency among T2 plants due to embryo or seedling lethality. Consequently, we analyzed these transformants to gain insights into the failure to establish *pigs-1/pigs-1, pPIGS:PIGS-GFP* plants among the progeny of *pPIGS:PIGS-GFP* complementation lines.

Aborted seeds caused by embryo lethality is one reason for distorted segregation of certain genotypes in progeny and can be readily scored in selfed pistils because they typically develop into wrinkled brown seeds, as reported in *pnt1* mutant that is defective in GPI biosynthesis (Gillmor et al., 2005). No shriveled or wrinkled seeds were observed in any of the three independent complementing *pigs-1/+, pPIGS:PIGS-GFP* lines. Additionally, immature siliques from line #15 contained similar number of unfertilized, aborted, and normally developed seeds as in wild type (Additional file 4), suggesting that absence of viable *pigs-1/pigs-1, pPIGS:PIGS-GFP* plants is not due to embryo lethality.

We next performed two experiments to test the possibility that *pigs-1/pigs-1, pPIGS:PIGS-GFP* plants died at the seedling stage after apparently normal looking seeds germinated. First, selfed seeds from *pigs-1/+, pPIGS:PIGS-GFP* plants were grown without selection on regular MS plates (containing no antibiotic or basta to select for the complementing *pPIGS:PIGS-GFP* transgene or the *pigs-1* mutation, respectively). We found several seedlings were stunted in growth and died shortly after emerging from the seed coat (% lethality ranging from 18-29; Table 3). Presence of lethal seedlings was dependent on the presence of *pPIGS:PIGS-GFP* transgene, as there were no lethal seedlings in the progeny of *pigs-1/*+, which lacked the complementing transgene (Table 3). Second, when selfed seeds from *pigs-1/+, pPIGS:PIGS-GFP* plants, but not when selfed seeds from *pigs-1/*+ plants, were plated on MS plates containing basta (marker to select *pigs-1* mutation), besides basta-resistant and basta-susceptible seedlings, we also detected lethal seedlings that died after emerging from the seed coat (Fig. 3a-c and Table 4). Importantly, lethal seedlings observed on MS basta plates were similar to those observed on regular MS plates, as they did not produce true leaves, failed to develop roots, and embryonic leaves appeared pale in color (Fig. 3c). These results suggested that *pigs-1/pigs-1* seeds were established in the presence of *pPIGS:PIGS-GFP* transgene but died after emerging from the seed coat.

**Figure 3.**
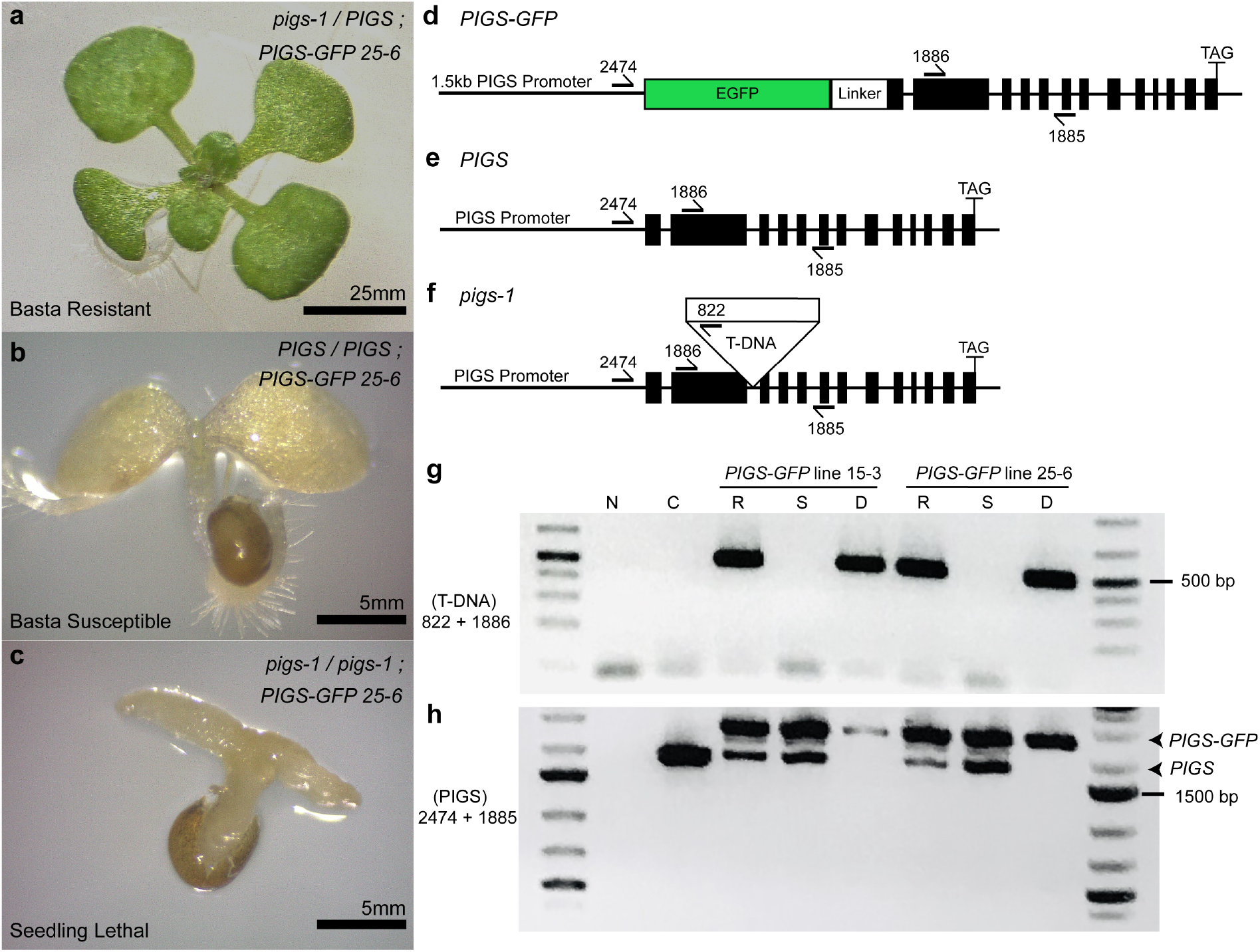
Lethal seedlings segregated in the progeny of selfed *pigs-1/+, pPIGS:PIGS-GFP*. (**a**-**c**) Light micrographs of seedlings from selfed seeds of *pigs-1/+, pPIGS:PIGS-GFP* line #25-6 grown on MS plates supplemented with basta for 14 days in constant light. (**a**) Basta-resistant *pigs-1/+, pPIGS:PIGS-GFP* seedlings had well developed true green leaves and prolific root, root hairs, and lateral root growth. (**b**) Basta-susceptible *+/+, pPIGS:PIGS-GFP* seedlings had minimal to no true leaves or root hair and lateral root development. (**c**) Lethal *pigs-1/pigs-1, pPIGS-PIGS-GFP* seedlings were small, showed aberrant pale cotyledon growth, and lacked radicle or roots. *Images shown were taken using a Zeiss Axiovert 100 microscope with either an 8x objective, Bar = 5mm (**a**) or 50x objective, Bar = 5mm (**b**-**c**). (**d**-**f**) Gene model of the (**d**) *pPIGS:PIGS-GFP* construct, (**e**) endogenous *AtPIG-S* gene, and (**f**) *pigs-1* T-DNA insertion. Primers used in the PCR-based genotyping of each of these three DNAs are indicated. (**g-h**) PCR-based genotyping using gDNA isolated from basta-resistant, basta-susceptible, and lethal seedling progeny of *PIGS-GFP* T2 lines #15-3 and #26-6 for either the presence of the *pigs-1* T-DNA using primers 822 + 1886 (**g**) or for the presence of the endogenous *AtPIG-S* or *PIGS-GFP* gene using primers 2474 + 1885. Expected size for endogenous *AtPIG-S* is 1896 bp and expected size for *PIGS-GFP* is 2586 bp. Original, un-cropped gels shown is provided in Additional file 5. N, water control; C, Columbia-0 gDNA; R, basta-resistant; S, basta susceptible; D, dead (lethal) seedlings. *All primer sequences listed in Additional Table 1.

**Table 4.**
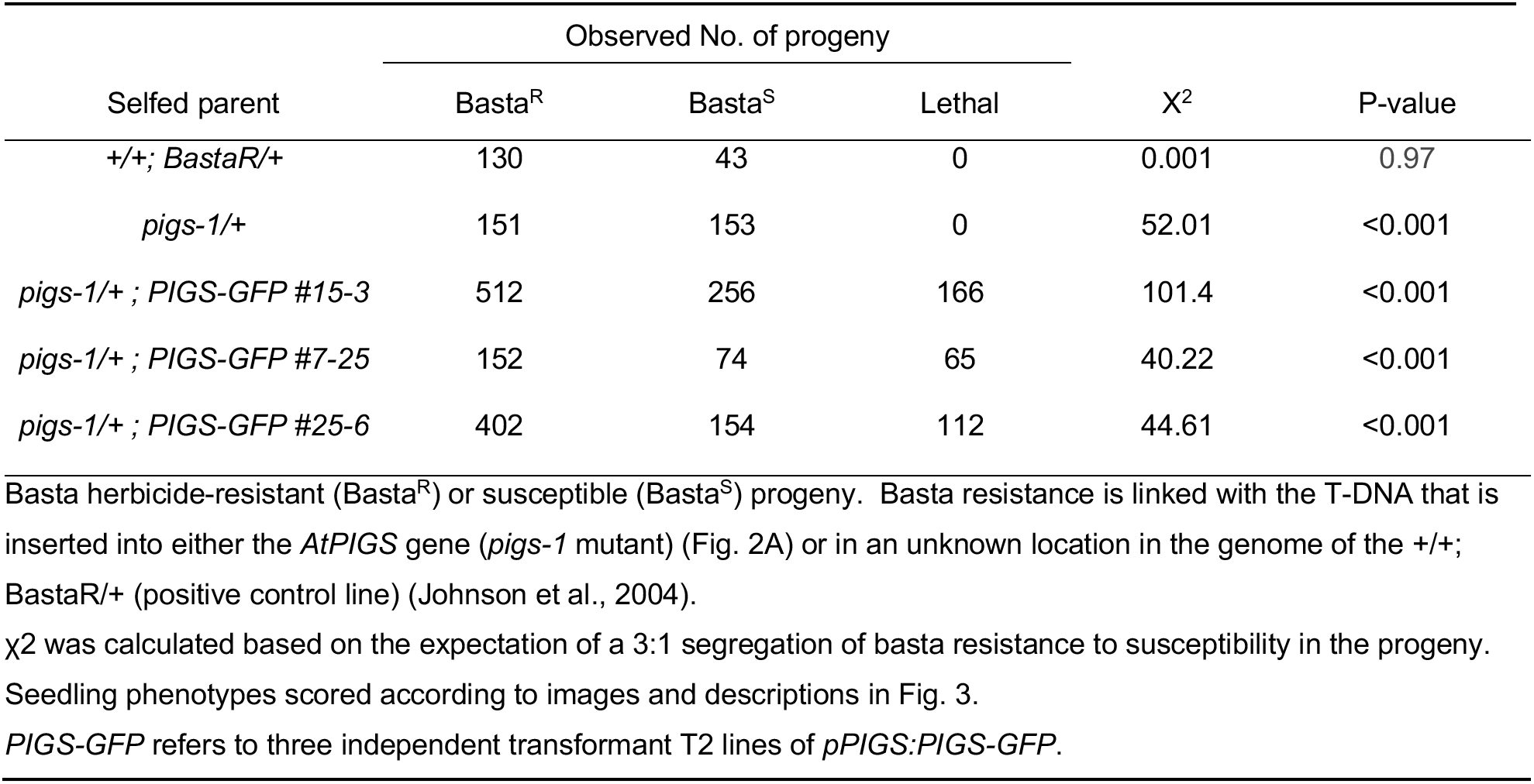
Transmission of *pigs-1* mutation in self-pollinated pistils.

We devised a PCR-based genotyping assay to verify if the lethal seedlings’ genotype was indeed *pigs-1/pigs-1, pPIGS:PIGS-GFP* (Fig. 3d-h and Additional file 5). For this, we scored the three types of seedlings in the progeny of two independent *pPIGS:PIGS-GFP* transformants grown on MS plates with basta, pooled each type of seedlings, isolated genomic DNA from them, and performed two sets of PCR reactions, one to score presence of the *pigs-1* mutation and the other for presence of the *pPIGS:PIGS-GFP* transgene and wild-type copy of *AtPIG-S* (Fig. 3d-h and Additional file 5). We found that only the pooled lethal seedlings, but not basta-resistant or basta-susceptible seedlings, were homozygous for the *pigs-1* mutation, confirming the genotype of lethal seedlings is indeed *pigs-1/pigs-1, pPIGS:PIGS-GFP* (Fig. 3g and h and Additional file 5). Care was taken to prevent seed coat contamination during seedling collection so that the maternal diploid genotype of the seed coat (*pigs-1/+, PIGS-GFP*) did not confound the PCR-based genotyping of seedlngs. Indeed, lack of endogenous wild-type *AtPIG-S* band in the lethal seedling samples confirmed that our seedling collection of this sample was free of seed coat contamination (Fig. 3h and Additional file 5). Additionally, the PCR results showed that the *pPIGS:PIGS-GFP* transgene was segregating in both pooled basta-resistant and basta-susceptible seedlings, and that pooled basta-susceptible seedlings did not contain the *pigs-1* mutation as expected (Fig. 3g and h and Additional file 5). These results showed that although the *pPIGS:PIGS-GFP* construct could rescue the pollen tube growth defects of *pigs-1* mutant pollen and allowed homozygous *pigs-1/pigs-1* seeds to be formed, the *AtPIGS-GFP* expressed from native *PIGS* promoter couldn’t complement the developmental defects in *pigs-1/pigs-1* seedlings.

## DISCUSSION

### Deficiency in GPI-anchoring of GAPs does not appear to disrupt the FG functions

Like that seen in *cis* mutations in the GPI-anchor signal sequence (GAS) of LRE (a substrate of GPI-T) and in *gpi8-2* (a component of GPI-T), *pigs-1* also disrupts the polar LRE-cYFP localization in the FA without affecting its own transmission through the FG. The GPI anchor of ZERZAUST, another Arabidopsis GAP, is dispensable for its biological activity (Vaddepalli et al., 2017), so it’s possible that LRE could simply be another GAP whose GPI anchor doesn’t influence its function. If so, the GAS in LRE could simply be a relic of its duplication where the ancestral copy contained a more functionally relevant GAS.

Still, presumably in *pigs-1* FGs, all GAPs lack GPI anchors, so it is unlikely that GPI anchor addition is unimportant for the function of all FG-expressed GAPs. One possibility is that retention of mature GAPs from the diploid *pigs-1/*+ megaspore mother cell (MMC), after meiosis and the three mitotic cell divisions in megagametogenesis, could support the function of the mature *pigs-1* FG. In support of this hypothesis, sporulating diploid yeast cells with a disruption in the *GPI2* gene produce GPI-deficient haploid ascospores that are able to germinate and complete up to four mitotic cell divisions (Leidich et al., 1995). Another possibility is that mature GAPs could be supplied by integument tissues that surround the FG, as it is known ‘membrane painting’ allows transfer of GAPs between cell surfaces of animal cells upon contact (Medof et al., 1996, Medof et al., 1984). Consistent with either of these possibilities, several GAPs involved in FG development such as AGP4, AGP18, A36, and A39 have mutant defects that are sporophytic in nature (Pereira et al., 2016, Acosta-Garcia and Vielle-Calzada, 2004, Gao et al., 2017).

Still, GAPs involved in pollen tube reception such as LRE and ENODLs are gametophytic in nature (Hou et al., 2016) and so loading from sporophytic MMC or integument tissues cannot completely explain why GAP-deficient FGs could still function normally. Interestingly, both LRE and ENODLs function in synergid cells, a polarized cell whose default secretory pathway is to direct protein traffic to the FA (Punwani et al., 2007). The high density of cell wall and membrane invaginations of the FA combined with the hydrophobicity of the un-cleaved GAS region of GAPs in *pigs-1* synergids could facilitate their sequestration in the FA (Galian et al., 2012). The synergid-secreted GAPs, ENODLs and LRE, could then be further retain at the *pigs-1* FA by virtue of their binding to ectodomains of receptor like kinases, such as FER (Liu et al., 2016, Li et al., 2015).

Finally, while *gpi8-2* and *pigs-1* mutations disrupt the polar localization of LRE-cYFP in the FA, some LRE-cYFP molecules still reach the FA of *pigs-1* (Figs. 2a-g) and *gpi8-2* (Liu et al., 2016) synergids. GAPs have the capacity to function at very low concentrations due to the high membrane dynamics of GPI anchors and their ability to rapidly “patrol” membranes (Medof et al., 1987, Rosse, 1990). Therefore, a 50% decrease in LRE-cYFP localization to the FA may not be sufficient to negatively impact its ability to function. In summary, the unusual structural and functional properties of synergids may overcome GPI anchor addition deficiencies to synergid cell-expressed GAPs and confound genetic analysis of GAPs in synergid cells.

### Loss of AtPIG-S function led to defects in pollen tube emergence and tube growth

Contrary to *pigs-1* mutant FGs, *pigs-1* mutant MGs have severe fertility defects. Here, by utilizing the GUS reporter within the *pigs-1* T-DNA we show for the first time that loss of AtPIG-S function in pollen (and by inference GPI transamidation-deficient pollen) have defects at the stages of pollen tube emergence and growth. Future investigations could focus on testing if low TE reported previously in *gpi8-1* and *gpi8-2* MGs are also caused by defects in pollen tube emergence and tube growth. It’s possible that loss of GPI anchor addition to pollen-expressed GAPs such as *LLG2* and *LLG3* (paralogs of LRE), which are critical for maintaining the cell wall integrity of pollen tubes, could be less tolerated in pollen than the FG because of the rapid developmental process pollen must undergo during tube emergence and growth (Ge et al., 2019). While experiments on localization of LLG2/3 in *pigs-1* pollen tubes would be difficult due to the low level of *pigs-1* pollen tube growth, analyzing the effect of *cis* mutations in the GAS region of LLG2/3, similar to those reported in LRE (Liu et al., 2016), could substantiate our findings on the increased importance of GPI anchor addition to GAPs in the MG compared to that in the FG.

Our results on *pigs-1* (this study) and *gpi8-2* (Liu et al., 2016) causing severe MG defects are also consistent with previous reports on mutations in four GPI biosynthesis genes causing a decrease in male fertility (Lalanne et al., 2004, Gillmor et al., 2005, Dai et al., 2014). While the mutant analyses on all four of these GPI biosynthesis genes established an important role of GPI in MG function, they could not preclude the possibility that the absence of free, non-protein linked GPI underlies these MG defects. In addition to serving as a substrate in the transamidation reaction catalyzed by GPI-T, evidence in protists suggests that free GPI plays an independent and more essential role in cell expansion than mature GAPs (Ilgoutz et al., 1999, Hilley et al., 2000). Here, by targeting the downstream transamidation step rather than GPI anchor biosynthesis, our study provides support to the possibility that the pollen tube emergence and growth defects in *pigs-1* pollen grains are due to a lack of GPI anchor attachment to GAPs. Still, free GPI may be important in pollen, but to our knowledge plasma membrane-localized free GPI in pollen has not been documented. Because *pigs-1* pollen grains are viable until tube emergence, they also offer a potential *in vivo* system to study free GPI in plants, as free GPI likely accumulates in *pigs-1* pollen grains, similar to GPI transamidation-deficient mutants in yeast (Meyer et al., 2000).

### A new role for *AtPIG-S* in Arabidopsis early stage seedlings is revealed by rescue of *AtPIG-S* function in the male gametophyte

Successful restoration of the *pigs-1* pollen tube growth defects by the *pPIGS:PIGS-GFP* transgene confirmed the role of *AtPIG-S* in the MG and suggested that fusion of GFP to the AtPIG-S protein did not affect its function in the GPI-T complex. Hence, *AtPIGS-GFP* fusion protein could be used to immunoprecipitate and unravel the subunit composition of the GPI-T complex in Arabidopsis similar to HA-PIG-S, which was used to pull down the GPI-T complex in CHO cells (Hong et al., 2003). However, it should be noted that in yeast, the *PIG-S* ortholog, GPI17, neither co-purifies (Fraering et al., 2001) nor stably interacts with other subunits of GPI-T (Zhu et al., 2005).

Rescue of AtPIG-S function in the MG by the *pPIGS:GFP-PIGS* transgene also allowed us to establish *pigs-1/pigs-1* seedlings. Like seedlings with a mutation in *PEANUT1 (PNT1)*, a predicted mannosyltransferase required for GPI biosynthesis (Gillmor et al., 2005), we showed that *pigs-1/pigs-1* seedlings carrying the *pPIGS:GFP-PIGS* transgene emerged from the seed coat but soon became necrotic. However, unlike *pnt1* mutant, the *pigs-1/pigs-1* seeds carrying the *pPIGS:GFP-PIGS* transgene were not wrinkled and shriveled. Previously, a role for GPI transamidation in seedling development and later stage plant development was proposed after characterizing the nonlethal allele, *gpi8-1*, a missense mutation in *AtGPI8* (Bundy et al., 2016). However, the aboveground organs of *gpi8-1* mutants were very minimally affected even 15 days after germination and produced roots, albeit significantly shorter than wild type. In contrast, *pigs-1/pigs-1, pPIGS:PIGS-GFP* seedlings died directly following emergence from the seed coat. Notably, the maternal seed coat is *pigs-1/*+ and perhaps haplo-sufficient to support the development of *pigs-1/pigs-1, pPIGS:PIGS-GFP* embryos. Still, it is not clear why the *pPIGS:PIGS-GFP* transgene failed to complement seedling function, though it was previously reported that in Arabidopsis, a *AtGPI8-EGFP* construct also driven under 2.1-kb of it’s endogenous promoter did not complement all *gpi8-1* phenotypes and that fluorescence reporter in this construct was not detectable (Bundy et al., 2016). It is possible that the promoter sequences used in our construct also lacked the *cis* elements required for sufficient expression in germinated seedlings, where *AtPIG-S* is expressed (http://bar.utoronto.ca/eplant/). Alternatively, the PIGS-GFP fusion protein could have failed to be recruited into the GPI-T complex specifically in seedlings. Nevertheless, by comparing *pnt1*, *pigs-1/pigs-1, pPIGS:GFP-PIGS,* and *gpi8-*1 phenotypes, it appears that seedling lethality in *pigs-1/pigs-1, pPIGS:GFP-PIGS* manifested at a developmental stage later than *pnt1* (Gillmor et al., 2005) but earlier than *gpi8-1* (Bundy et al., 2016). Therefore, our complementation lines can be used to dissect the role of GPI anchor addition to GAPs and the presence of AtPIG-S in the GPI-T complex in early stage Arabidopsis seedlings.

## CONCLUSIONS

During flowering plant reproduction, GAPs mediate many MG and FG functions. While the loss of GPI anchor addition to the GAPs did not result in noticeable phenotype(s) in the FG, its importance in the MG is underscored by the pollen tube emergence and growth defects of *pigs-1* mutant pollen. Up to this point, MG defects in GPI anchor biosynthesis mutants could not preclude the possibility that the absence of free, non-protein linked GPI underlie their MG defects. Here, instead we analyzed a putative subunit in the GPI-T complex, which is involved in post-translational addition of GPI anchors to GAPs, and show that MG defects of GPI-deficient pollen grains seen previously are likely due to a lack of mature GAPs.

We also substantiate the finding that loss of GPI anchor addition to the GAPs in the FG does not affect its functions. While it’s possible the megaspore mother cell could have supplied mature GAPs to the FG across cell divisions, the pollen tube reception defects seen in the FG-expressed GAPs such as LRE and ENODLs are gametophytic in origin. Therefore, we propose that the importance of their GPI anchors in the FG-expressed GAPs could be masked by their synergid expression such that (1) the specialized secretion system of synergids is sufficient to transport non-GPI anchored GAPs to the plasma membrane, and (2) the high surface area of the FA in combination with the hydrophobicity of their un-cleaved GAS region are sufficient for their sequestration and function at the FA. For these reasons, the MG, compared to FG, is a better haploid model to examine the role of GAPs.

## METHODS

### Plant materials and Growth conditions

Columbia (Col-0) is the ecotype of all Arabidopsis seeds used in this study. The *gpi8-2/*+ (CS853564), *pigs-1/*+ (CS807841), and *pigt-1* (Salk_099158) seeds were purchased from Arabidopsis Biological Resource center (ABRC) in Columbus, Ohio. The *pLAT52:GUS* gene in the wild-type background was described previously (Johnson et al., 2004) and is a donation from Dr. Mark Johnson, Brown University, Providence, RI, USA. Once the manuscript has been published, newly established transgenic lines generated as part of this study – *pLRE:LRE-cYFP/pLRE:LRE-cYFP, pigt-1/pigt-1*; *pLRE:LRE-cYFP/pLRE:LRE-cYFP, pigs-1/+*, and *pPIGS:PIGS-GFP, pigs-1/+* – by Nick Desnoyer, the first author of this study, will be deposited in the Arabidopsis Biological Resource Center in Columbus, Ohio, USA. All Arabidopsis plants reported in this study were grown as described (Tsukamoto et al., 2010).

### Pollen tube growth assays

*In vivo* pollen tube growth assays were done as described (Tsukamoto et al., 2010). Briefly, *pLAT52:GUS* or *pigs-*1/+ pollen was crossed to emasculated stage 14 pistils of the indicated plants. Crossed pistils were then collected 18 hours after pollination, fixed in 80% acetone, stained for GUS activity using x-gluc (Johnson et al., 2004), mounted in 50% glycerol, and imaged using differential interference contrast optics in a Zeiss Axiovert 100 microscope. Light micrographs of pistils were used to measure pollen tube length using ImageJ.

### Transmission efficiency assay

Reciprocal crosses were done on emasculated stage 14 pistils. For limited pollination experiments, less than 40 pollen grains were used to pollinate pistils under a dissecting microscope. Seeds from these crosses were placed on plates containing appropriate antibiotics/herbicides corresponding to the antibiotic or herbicide resistance marker in the T-DNA. 14-day old seedlings with true leaves and developing roots were scored as resistant.

### Seedling lethal assay

Seeds were plated on either MS plates with or without basta, stratified for 2 days at 4 °C, incubated in the growth chamber for 14 days and scored under a dissecting scope. In basta plates, three types of seedlings were scored. If seedlings contained true leaves and all leaves were green, and had robust root growth with root hairs and secondary roots, they were scored as ‘basta-resistant’. If the germinated seedlings contained fully developed embryonic leaves but true leaves never emerged, and radicle and roots emerged but failed to grow and after two weeks the seedling died, they were scored as ‘basta-susceptible’. Finally, if seedlings that began to emerge from the seed coat contained only embryonic leaves and contained neither radicle nor roots, they were scored as ‘lethal seedlings’.

In case of seeds plated on MS plates without basta, two types of seedlings were scored; if seedlings contained true leaves and all leaves were green, and had robust root growth with root hairs and secondary roots, they were scored as ‘normal’ and if seedlings that began to emerge from the seed coat contained only embryonic leaves and contained neither radicle nor roots, they were scored as ‘lethal’ seedlings.

For genomic DNA isolation, seedlings were collected after 14 days of growth, taking care not to contaminate the collection with seed coat, especially in case of lethal seedlings. Although lethal seedlings were detected within 7-10 days after plating, we scored and collected the three types of seedlings after 14 days of growth so that it was possible to separate and collect lethal seedlings without seed coat sticking to the seedling and contaminating the collection. Avoiding seed coat contimation was essential so that diploid maternal genotype of the seed coat did not confund the PCR genotyping of seedlings. Each of the three type of seedlings were collected separately, pooled and frozen in liquid nitrogen, and genomic DNA was isolated from them as described previously (Tsukamoto et al., 2010). PCR was performed with at least 100ng of genomic DNA as a template in each reaction as follows: Step 1: (1X) 98.0 °C for 2 minutes; Step 2: (34X) 95.0 °C for 30 seconds, 56.0 °C for 20 seconds, and 72.0 °C for 1 minute; Step 3: (1X): 72.0 °C for 2 minutes. PCR products were separated on a 1% agarose gel. Sequence of primers used and expected size of PCR products are listed in Additional Table 1.

### Confocal Microscopy

The *pigt-1/pigt-1* or *pigs-1/*+ mutant was crossed to *pLRE:LRE-cYFP/pLRE:LRE-cYFP* (Liu et al., 2016) and established *pigt-1/pigt-1*, *pLRE:LRE-cYFP/pLRE:LRE-cYFP* and *pigs-1/*+, *pLRE:LRE-cYFP/pLRE:LRE-cYFP* lines, respectively. Confocal images of ovules from these lines were taken after mounting unpollinated stage 14 pistils. Fluorescent images were taken using a Leica SP5 confocal laser scanning microscope system. For cYFP imaging, samples were excited with a 488-nm laser, and emission spectra between 510 and 550 nm were collected. YFP images were processed with ImageJ software (http://imagej.nih.gov/ij/). YFP localization was quantified by measuring raw integrated density using the ROI manager tool in ImageJ.

### RNA isolation, RT PCR, and RT-qPCR

RNA was isolated according to (Qin et al., 2009). Briefly, for each biological replicate, 14 days-old *pigt-1/pigt-1* and Col-0 seedlings were collected, flash frozen, and stored at −80°C until RNA extraction. Three biological replicates were collected for both genotypes. RNA was isolated using RNeasy Plant Mini Kit (QIAGEN, Catalog # 74904) and subjected to RNase-free DNase I (Life Technologies, Catalog # AM2222) treatment to remove potential DNA contamination, cleaned up using RNeasy MinElute Cleanup Kit (QIAGEN, Catalog # 74204), and tested for RNA integrity using an Agilent Bioanalyzer 2100 (Agilent Technologies, Boblingen, Germany). For RT-PCR, cDNA was synthesized from 2 μg total RNA using SuperScript™ IV First-Strand Synthesis System, ThermoFisher Scientific, Catalog #18091050.

*PIGT* PCR on *pigt-1/pigt-1* and Col-0 cDNA was amplified using either primer pairs (P1+P2) or (P1+P4) with the following program: Step 1 (1X): 98.0 °C for 2 minutes; Step 2 (37X): 98.0 °C for 15 seconds, 55.0 °C for 15 seconds, and 72.0 °C for 2 minutes. Sequence of primers used and expected size of PCR products are listed in Additional Table 1.

The following qPCR program was used for all qPCR experiments: Step 1 (1X): 95.0 °C for 10 minutes; Step 2: (40X) 95.0 °C for 10 minutes, 55.0 °C for 30 seconds and 72.0 °C for 30 seconds, Data collection and real-time analysis enabled; Step 3 (101X): 45.0 °C-95.0 °C for 10 seconds, increase set point temperature after Step 2 by 0.5 °C, Melt curve data collection and analysis enabled. Ct values were normalized to *ACTIN2/8*. Relative levels of gene expression were calculated according to Qin et al., 2009. At least two technical replicates of qPCR were performed for each experiment. Primers used in qPCR reactions are listed in Additional Table 1.

### Cloning *pPIGS:GFP-PIGS*

The *pPIGS:GFP-PIGS* construct was created by overlap PCR with PrimeSTAR® GXL DNA Polymerase (TaKaRa Bio Inc.; Catalog # R050A) and DNA templates and primers listed in Additional Table 1 and cloned into pH7WG plasmid linearized with SalI-HF (NEB, Cataog # R3138S) and AscI (NEB, Catalog # R0558S) by using the In-Fusion HD Cloning Plus system (Clontech, Catalog # 639645). The recombinant plasmids were transformed into Stellar™ Competent Cells (Clontech, Catalog # 636763), and positive colonies were selected on LB plates containing spectinomycin (100ug/mL, Sigma-Aldrich, Catalog # 85555). The construct was sequence verified (Eton Bioscience, Inc.) before transforming into *Agrobacterium tumefaciens* (GV3101 pMP90 strain). The positive colony selected for transforming into Arabidopsis was also verified by colony PCR for the presence of the transgene.

### Plant Transformation

*pigs-1/*+ heterozygous inflorescences were dipped into transformation solution containing *Agrobacterium tumefaciens* (GV3101 pMP90 strain) harboring the *pPIGS:GFP-PIGS* plasmid (Chung et al., 2000). Hygromycin-resistant transformants were selected as described (Harrison et al., 2006) with a Hygromycin concentration of 20ug/mL. T1 seeds were plated and stratified for 2-3 days, placed in a plant growth chamber (set at 21°C with continuous light (PHILIPS F17T8/TL741 Fluorescent Tube Light Bulb, 75-100 μmol·m^−2^·s^−1^) for 5-6 hours, then moved to darkness at room temperature for 3-4 days. Plates were then placed back into the growth chamber and transformants were selected based on presence of true leaves, which were present only in hygromycin-resistant plants.

### Transient expression of *pPIGS:GFP-PIGS* in *Nicotiana benthamiana*

An *Agrobacterium* GV3101::pMP90 strain carrying the plasmid containing *pPIGS:GFP-PIGS* was used to transiently express GFP-PIGS fusion protein in *N. benthamiana* as described (Zhan et al., 2018). A leaf punch was taken 3 days or 12 days post-infiltration from the infiltrated area and mounted onto 50% glycerol. The bottom epidermis of the leaf disk was then analyzed by confocal microscopy. Images were taken using a 488nm wavelength at 10.5% laser power on a Zeiss LSM 880 AxioObserver.

### Image Processing

Photoshop CC 2018, Illustrator CC 2019 (Adobe, https://www.adobe.com/) and ImageJ were used for assembling image panels and preparing figures.

### Accession Numbers

Accession numbers of genes studied in this work: *LRE* (*At4g26466*), *AtGPI8 (At1G08750),* and *AtPIGT (At3G07140)*.

## Supplementary information

**Additional file 1.** *pigt-1* mutation does not affect polar localization of LRE-cYFP in the filiform apparatus of synergid cells.

**Additional file 2**. Analysis of *PIG-T* expression in the *pigt-1* mutant.

**Additional file 3.** Transient expression of AtPIGS-GFP protein in *Nicotiana benthamiana* leaves.

**Additional file 4.** Selfed seed set in T2 *pigs-1/+, pPIGS:PIGS-GFP* plants.

**Additional file 5.** PCR-based genotyping of *pigs-1/pigs-1, pPIGS:PIGS-GFP* plants.

**Additional Table 1.** List of primers used in this study.

## Declarations

## Ethics approval and consent to participate

Not applicable

## Consent for publication

Not applicable

## Availability of data and materials

Data sharing not applicable to this article as no datasets were generated or analysed during the current study.

## Abbreviations

GPI: Glycosylphosphatidylinositol
GPI-T: GPI transamidase complex
GAPs: GPI anchors to GPI-anchored proteins
FG: Female Gametophyte
MG: Male Gametophyte
LRE: LORELEI
FA: Filiform Apparatus
FER: FERONIA
TE: Transmission Efficiency
MMC: Megaspore Mother Cell

## Authors’ contributions

N.D. and R.P. designed the experiments; N.D. established all three transgenic lines (*pLRE:LRE-cYFP/pLRE:LRE-cYFP, pigt-1/pigt-1*; *pLRE:LRE-cYFP/pLRE:LRE-cYFP, pigs-1/+*, and *pPIGS:PIGS-GFP, pigs-1/+*) generated newly as part of this study. N.D., G.H., and E.J. performed the experiments; N.D. and R.P. wrote the paper. All authors have read and approved the manuscript.

## Acknowledgements

This research was supported by a NSF grant to R.P. (IOS-1146090) and the Boynton Graduate Fellowship in Plant Molecular Biology, School of Plant Sciences, University of Arizona and University of Arizona Graduate Professional Student Council to N.D. We thank the Yadegari lab for help with transient expression of *pPIGS:GFP-PIGS* in *Nicotiana benthamiana*. We thank Patty Jansma in the Marley Imaging Core Facility for help with confocal microscopy.

