## Supplementary_Information for "AtPIG-S, a predicted Glycosylphosphatidylinositol Transamidase Subunit, is critical for pollen tube growth in Arabidopsis"

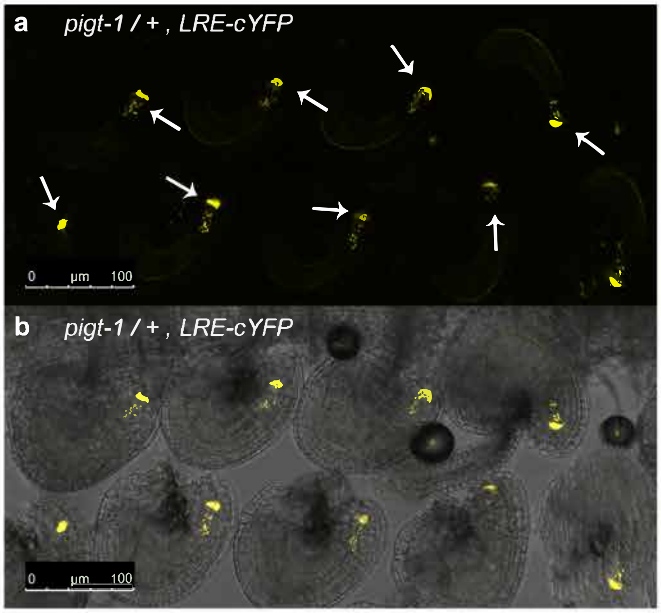


**Additional file 1**

**Additional file 1.** *pigt-1* mutation does not affect polar localization of LRE-cYFP in the filiform apparatus of synergid cells.

(**a**) and (**b**) Localization of the LRE-cYFP fusion protein in a *pigt-1/*+ pistil. **(a)** A representative fluorescent image of a portion of the pistil captured in the YFP channel of a confocal microscope showing LRE-cYFP localization. White arrows indicate synergid cells with a polarized cYFP localization in the filiform apparatus. **(b)** Merged view of fluorescent image in A with the bright field image of the same portion of the pistil captured in A. Bar = 100 μm.

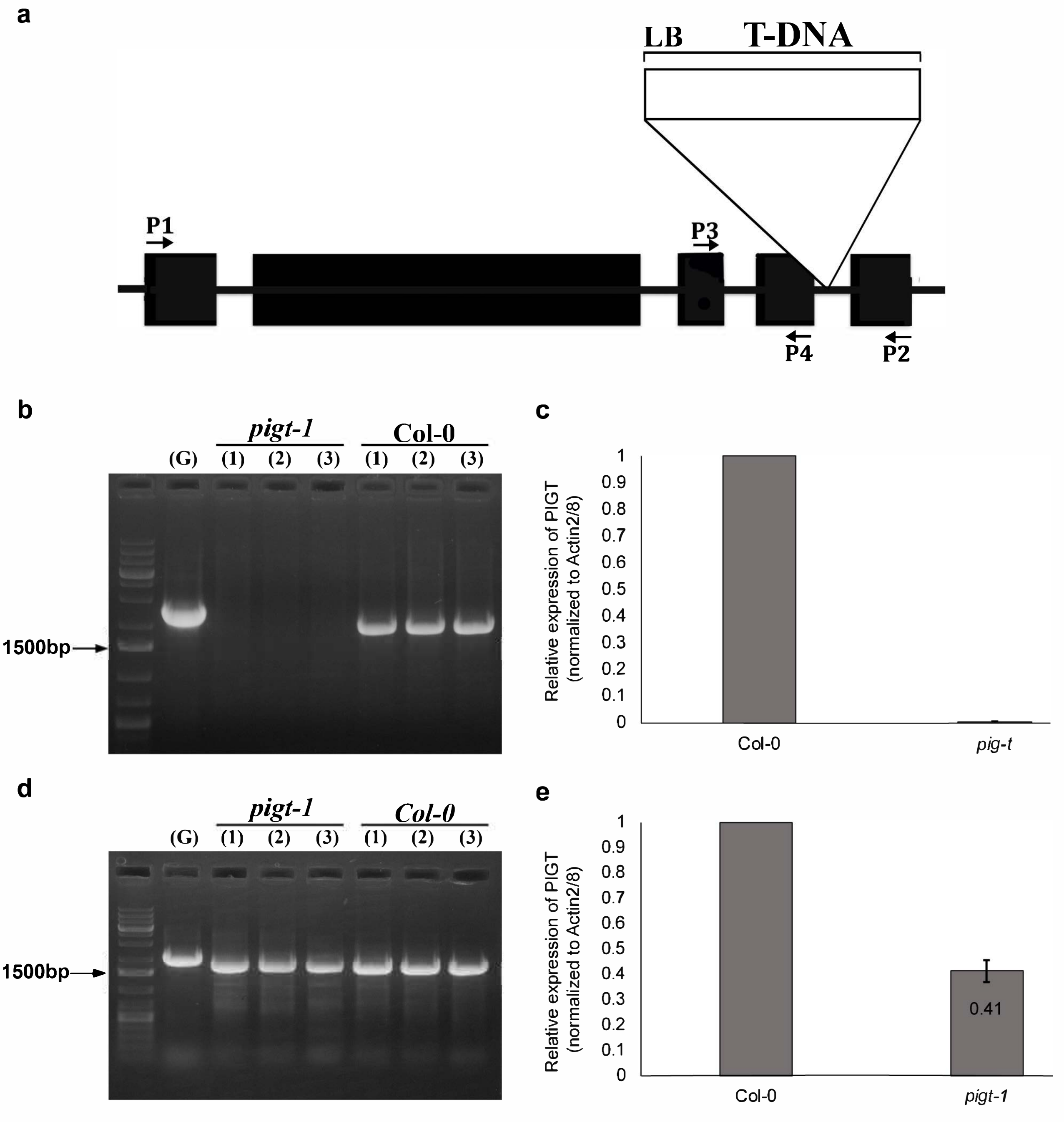


**Additional file 2**

**Additional file 2**. Analysis of *PIG-T* expression in the *pigt-1* mutant.

(**a**) Gene model of *AtPIG-T* (*AT3G07140*) with a T-DNA insertion (*Salk_099158*). Left border of T-DNA was sequenced showing an insertion in the fourth intron. Black rectangles indicate exons and lines indicate introns and UTRs. Primers used in RT-PCR experiments are indicated with arrows (P1-P4) and are found in Additional Table 1.

(**b**, **d**) RT-PCR analysis of either (**b**) full-length *AtPIG-T* using primers P1 and P2, or (**d**) partial length *AtPIG-T* exons 1 through 4 using primers P1 and P4 in cDNAs isolated from wild-type or homozygous *pigt-1* mutant 14-day old seedlings. G refers to genomic DNA used as template in the PCR reaction and #s 1-3 refer to three biological replicates of wild-type (Col-0) or *pigt-1* tissues.

(**c**, **e**) RT-qPCR analysis of either (**c**) *AtPIG-T* exons 3 through 5 using primers P2 and P3, or (**e**) *AtPIG-T* exons 3 through 4 using primers P3 and P4 in cDNAs isolated from wild-type or homozygous *pigt-1* mutant 14-day old seedlings. Ct values were normalized to *ACTIN2/8* and relative levels of gene expression were calculated by raising 2 to the power of (WT – *pigt-1* normalized Ct values) according to (Qin et al., 2009). Three biological replicates of wild type (Col-0) or *pigt-1* tissues and at least two technical replicates of qPCR were performed for each experiment.

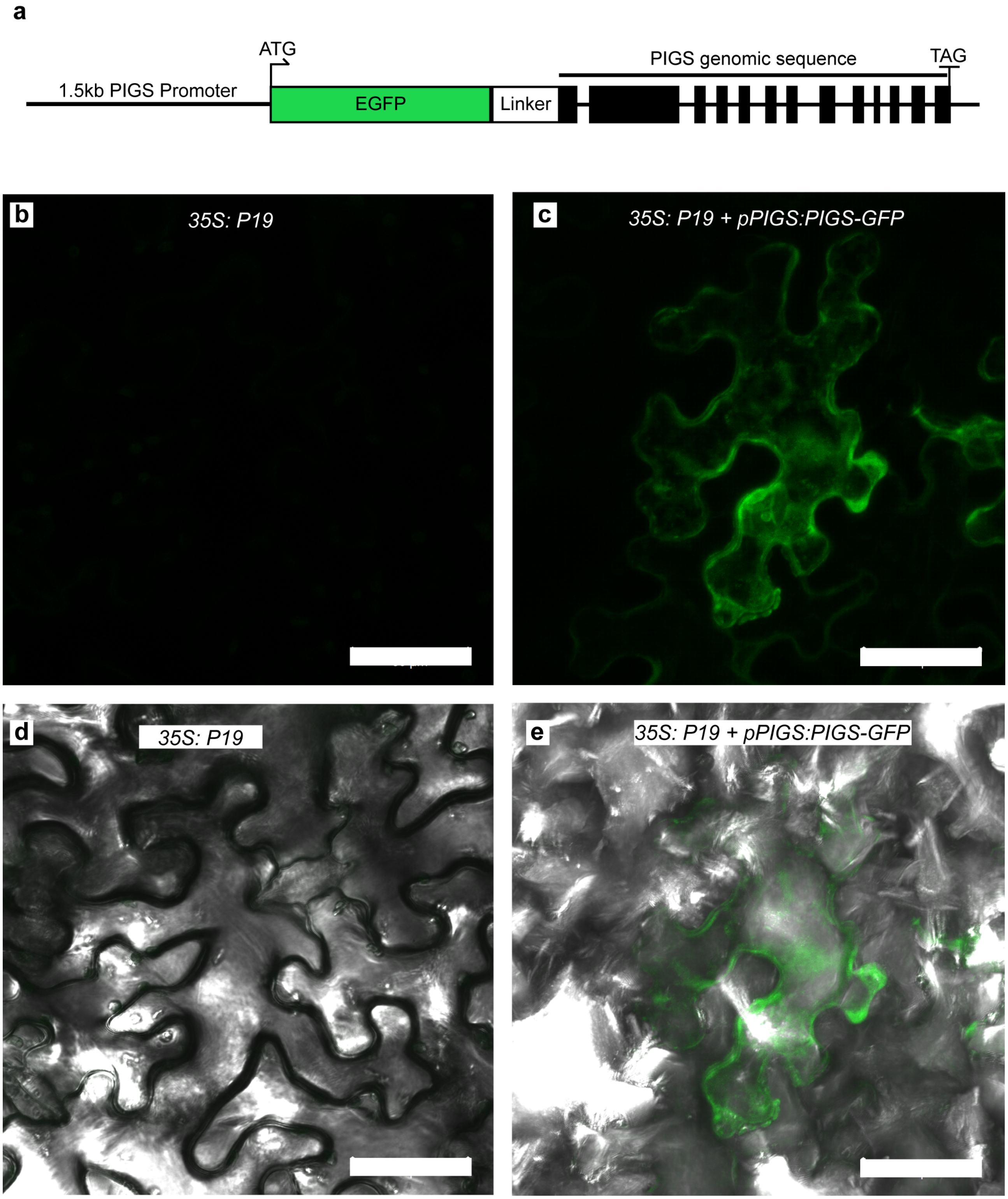


**Additional file 3**

**Additional file 3.** Transient expression of AtPIGS-GFP protein in *Nicotiana benthamiana* leaves.

(**a**) Gene model of *pPIGS:PIGS-GFP* construct. 1500 base pairs upstream of the *PIGS* transcriptional start site were fused to the translational start site of *EGFP* and linked to the At*PIG-S* genomic sequence by a glycine linker.

(**b**, **c**) Fluorescent images of *Nicotiana benthamiana* pavement cells 12 days post infiltration with either only *35S:P19* (helper plasmid) (**b**) or both *35S:P19* and *pPIGS:PIGS-GFP* (**c**).

(**d**, **e**) Merged view of fluorescent images in **b** and **c** with the bright field image of the same portion of the leaf.

Confocal images were taken at the same settings using a 488nm wavelength at 10.5% laser power on a Zeiss LSM 880 AxioObserver. Scale bars 50μM.

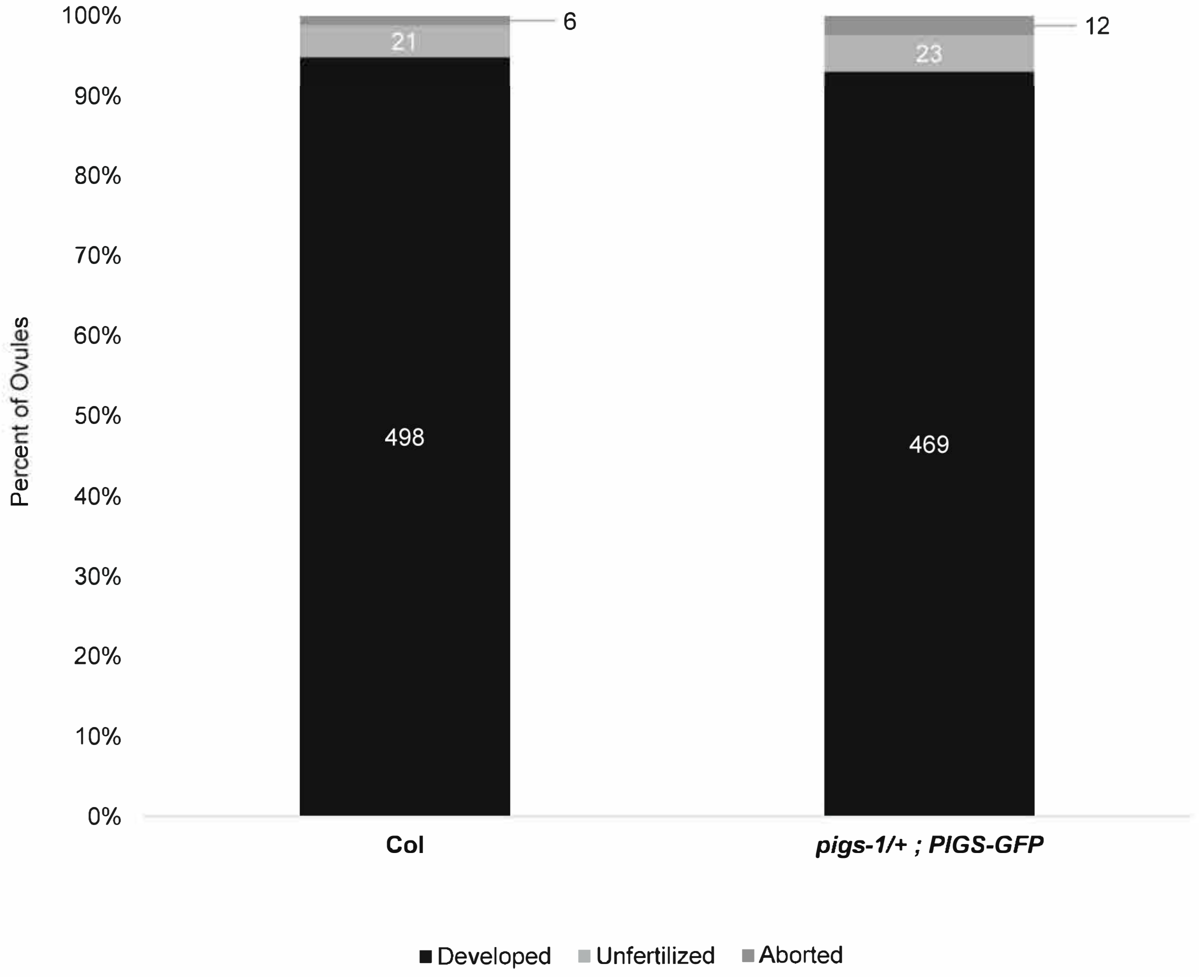


Additional file 4

**Additional file 4.** Selfed seed set in T2 *pigs-1/+ ; pPIGS:PIGS-GFP* plants.

Seeds within five indehiscent mature siliques from two wild-type and two *pigs-1/+* plants carrying at least one copy of the *pPIGS:PIGS-GFP* transgene were scored and categorized as either developed (normal seeds: black bar), aborted (pale color and collapsed seeds: dark gray bar), or unfertilized (shriveled and miniscule ovules: light gray bar). Numbers in the column refer to the number of seeds or ovules that were scored for each category.

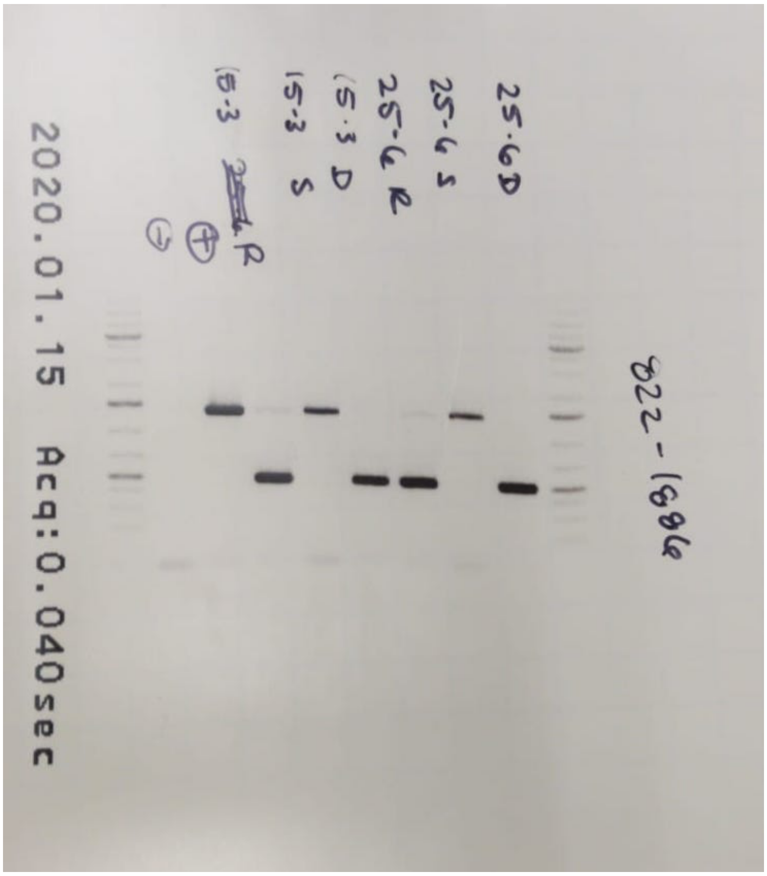


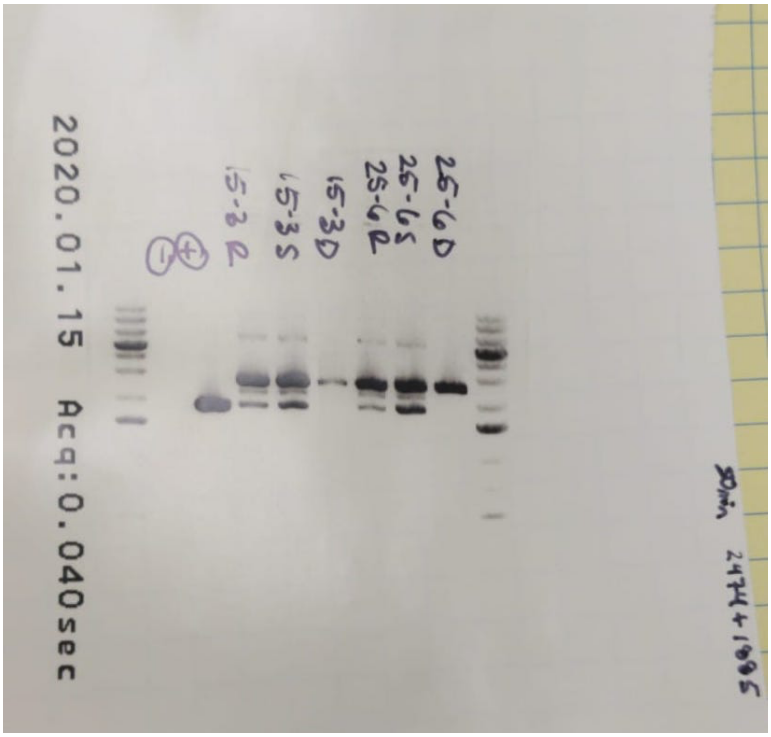


**Additional file 5**

**Additional file 5.** PCR-based genotyping of *pigs-1/pigs-1, pPIGS:PIGS-GFP* plants. Original, un-cropped gels of PCR reactions to genotype the presence of T-DNA insertion in *PIG-S* locus in *pigs-1*, endogenous *PIG-S* gene, and wild-type *PIG-S* gene in the *pPIGS:PIGS-GFP* transgene among the progeny of *pigs-1/+, pPIGS:PIGS-GFP* plants are shown here. Edited versions (for clarity) of these two images were provided in Fig. 3g, h.

**Additional Table 1. List of primers used in this study**

| Primer # | Primer Name | Sequence (5' - 3') | Template | Primer Pair | Expected Length (bp) |
|---|---|---|---|---|---|
| *pPIGS::GFP-PIGS* construct | | | | | |
| 2366 | PIGS Promoter_F | CCCATATGGTCGACCTGCCTCATGAATATCATTAA GATCCAGA | Col-0 genomic DNA | 2366 + 2368 | 1534 |
| 2368 | PIGS Promoter_R | CGCCCTTGCTCACCATCGGTGCTTTCCGAGAG | | | |
| 2367 | GFP_F | CTCTCGGAAAGCACCGATGGTGAGCAAGGGCG | pCAMBIA1300-GFP | 2367 + 2369 | 756 |
| 2369 | GFP_R | CCACCTCCACCTCCAGGCCGGCCCTTGTACAGCT CGTCCATGCCG | | | |
| 2370 | PIGS_F | GCCTGGAGGTGGAGGTGGAGCTATGGAAGAAAT CTCCGATCGT | Col-0 genomic DNA | 2370 + 2371 | 3280 |
| 2371 | PIGS 3'UTR_R | GTCGGCGCGCCCACCGGTTAGACATCTGTAATCG TAAAC | | | |
| *PIGT* RT-PCR | | | | | |
| 2364 | PIGT Full Length Forward (P1) | ATGGCTAGTCTTCTTCGATCC | Col-0 genomic DNA cDNA | 2364 + 2365 | Genomic: 2355 cDNA: 1935 |
| 2365 | PIGT Full Length Reverse (P2) | CTACTCGTCCGTGGAAAAATATTG | | | |
| 2364 | PIGT Full Length Forward (P1) | ATGGCTAGTCTTCTTCGATCC | Col-0 genomic DNA cDNA | 2364 + 2365 | Genomic: 2063 cDNA: 1747 |
| 2381 | PigT_exon4R (P4) | CTTGGCTTTTGAGAAACCTTTC | | | |
| 464 | ACTIN2/8_F | CCTATTGAGCATGGTGTTGTTAGCAAC | Col-0 genomic DNA cDNA | 464 + 465 | Genomic: 373 cDNA:277 |
| 465 | ACTIN2/8_R | TGTGAGACACACCATCACCAGA | | | |
| *PIGT* qRT-PCR | | | | | |
| 1890 | AT3G07140-1702F (P3) | CTTTGATAAGCTTCCCCGATC | cDNA | 1890 + 2365 | 452 |
| 2365 | PIGT Full Length Reverse (P2) | CTACTCGTCCGTGGAAAAATATTG | | | |
| 1890 | AT3G07140-1702F (P3) | CTTTGATAAGCTTCCCCGATC | cDNA | 1890 + 2381 | 264 |
| 2381 | PigT_exon4R (P4) | CTTGGCTTTTGAGAAACCTTTC | | | |
| 464 | ACTIN2/8_F | CCTATTGAGCATGGTGTTGTTAGCAAC | cDNA | 464 + 465 | 277 |
| 465 | ACTIN2/8_R | TGTGAGACACACCATCACCAGA | | | |
| *pigt-1* genotyping | | | | | |
| 1889 | AT3G07140-2720R (SALK_099158 LP) | GGATGCAACAAGAGAAAGCTG | *pigt-1* genomic DNA | 1889 + 1890 | 1019 |
| 1890 | AT3G07140-1702F (SALK_099158 RP) | CTTTGATAAGCTTCCCCGATC | | | |
| 383 | SALK-LB-6108R (pBIN-pROK2 LB) | CCAGCCAACAGCTCCCCGAC | *pigt-1* genomic DNA | 383 + 1890 | ~600 |
| 1890 | AT3G07140-1702F (SALK_099158 RP) | CTTTGATAAGCTTCCCCGATC | | | |

| | | | | | |
|---|---|---|---|---|---|
| *pigs-1* and *PIGS* genotyping | | | | | |
| 1885 | AT3G07180-1700R (SAIL_162_D06_LP) | GAAGGACATATTGACGCAAGG | *pigs-1* genomic DNA | 1885 + 1886 | 1042 |
| 1886 | AT3G07180-658F (SAIL_162_D06_RP) | TAAAGAGAATGCCAATGGTGG | | | |
| 822 | SAIL-LB-451R (pDAP101 LB) | GCCTTTTCAGAAATGGATAAATAGCCTTGCTTCC | *pigs-1* genomic DNA | 822 + 1886 | ~500 |
| 1886 | AT3G07180-658F (SAIL_162_D06_RP) | TAAAGAGAATGCCAATGGTGG | | | |
| 1885 | AT3G07180-1700R (SAIL_162_D06_LP) | GAAGGACATATTGACGCAAGG | *pigs-1* genomic DNA | 1885 + 2474 | PIGS-GFP: 2586 PIGS: 1896 |
| 2474 | PIGS Promoter | CGCTGAGCTAAGACGGCTATTG | | | |
